# NUAK1 regulates contraction in vascular smooth muscle cells derived from abdominal aortic aneurysm patients

**DOI:** 10.1101/2024.12.19.629332

**Authors:** Karlijn B. Rombouts, Tara A.R. van Merrienboer, Alex A. Henneman, Jaco C. Knol, Thang V. Pham, Sander R. Piersma, Connie R. Jimenez, Peter L. Hordijk, Jolanda van der Velden, Natalija Bogunovic, Kak Khee Yeung

## Abstract

**Aim:** Abdominal aortic aneurysms (AAA) are defined as a dilatation of the aortic wall. Impaired contractile ability of vascular smooth muscle cells (vSMC) is a hallmark of AAA. This study investigates the underlying mechanism of altered *in vitro* contractility of vSMC derived from AAA patients (AAA-SMC) compared to control vSMC (C-SMC).

**Methods:** Contractility of AAA-SMC (n=24) and C-SMC (n=8) was measured using Electric Cell-substrate Impedance Sensing. Large variability in AAA-SMC contraction was observed compared to C-SMC contraction, and AAA-SMC were therefore subdivided into low, intermediate (i.e. comparable to C-SMC contraction) and high contracting. To identify novel proteins involved in altered AAA-SMC contraction, a phosphoproteomic analysis was performed.

**Results:** The proteomics data showed that Thrombospondin-1, PDZ and LIM domain protein 4 and ATPase plasma membrane Ca^2+^ transporting 1 expression correlated with vSMC contraction, but knockdown (KD) of these targets did not affect contraction. Next, the phosphoproteomics data identified NUAK family kinase 1 (NUAK1) as a potential regulator of contraction, since its kinase activity correlated with AAA-SMC contraction. NUAK1 regulates Myosin phosphatase targeting subunit 1 (MYPT1) activity by phosphorylation, as confirmed by the correlation between NUAK1 activity and phosphorylation levels of Ser445 and Ser910 on MYPT1. NUAK1 protein and RNA expression correlated with AAA-SMC contraction. Moreover, NUAK1 KD decreased contraction in AAA-SMC, combined with reduced phosphorylation levels of Myosin light chain (pMLC) and Vinculin gene expression, and increased F-actin cytoskeletal filament levels.

**Conclusions:** NUAK1 regulates AAA-SMC contraction. Low NUAK1 expression decreased pMLC, potentially by higher MYPT1 activity, and subsequently reduced contraction.

**Graphical abstract:** 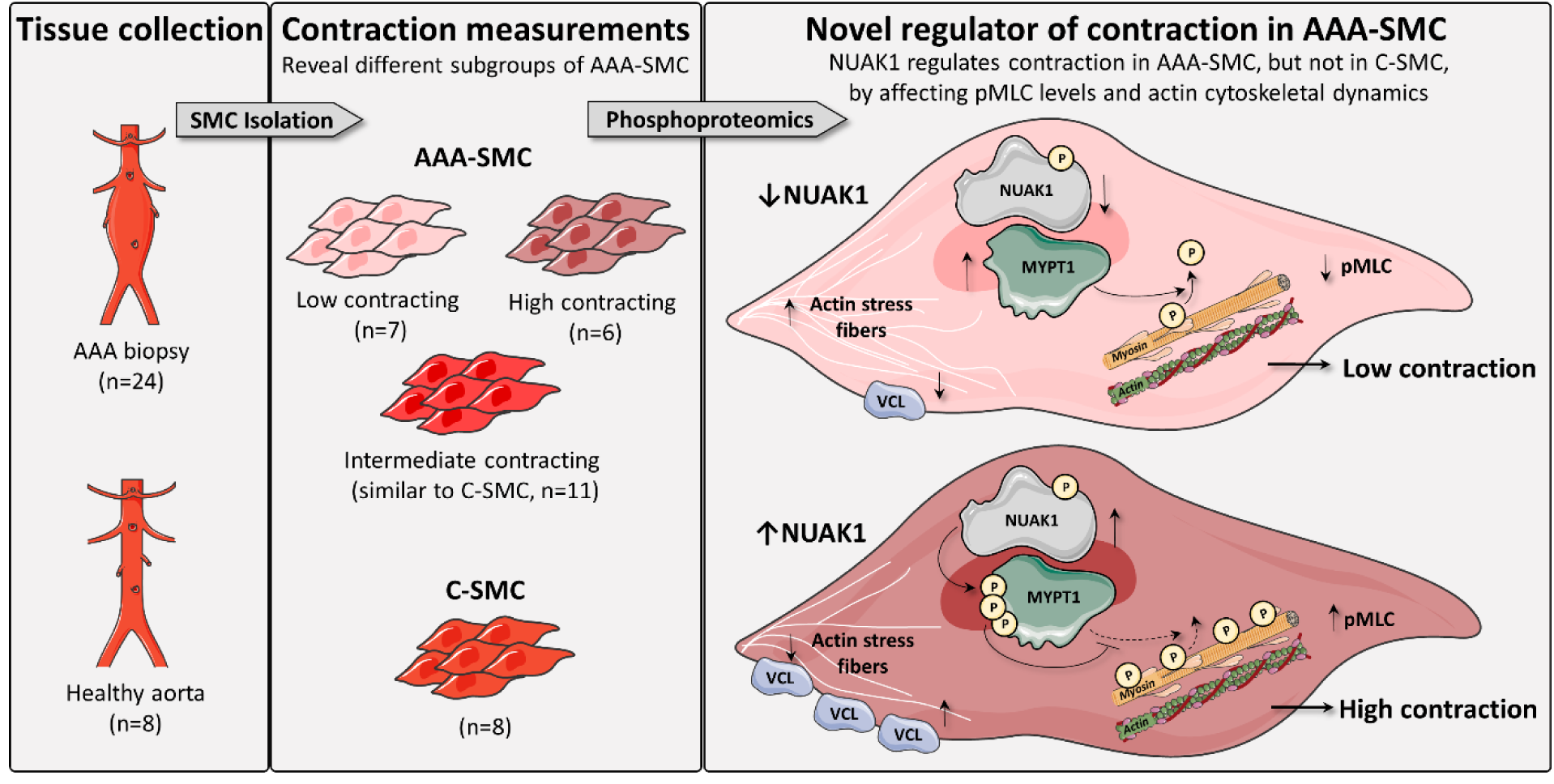

Elements were modified from Servier Medical Art, licensed under a Creative Common Attribution 3.0 Generic License. https://smart.servier.com/; https://creativecommons.org/licenses/by/3.0/

**Highlights:**

- Quantification of *in vitro* vascular smooth muscle cells (vSMC) contractile capacity revealed large variation in contraction of vSMC derived from human AAA tissue (AAA- SMC) compared to vSMC derived from healthy aortic biopsies (C-SMC).
- Phosphoproteomic analysis identified NUAK family kinase 1 (NUAK1) as a novel regulator of contraction in AAA-SMC, since its kinase activity and expression levels correlated with AAA-SMC contraction.
- NUAK1 regulates contraction in AAA-SMC by regulating Myosin phosphatase targeting subunit 1 (MYPT1) activity through phosphorylation, which subsequently affects phosphorylation levels of Myosin light chain (pMLC), and by affecting Vinculin expression and F-actin cytoskeletal filament levels.
- Gaining more insight into NUAK1 function and mechanisms leading to aortic wall weakening can ultimately contribute to better AAA disease prediction and treatment.

## 1. Introduction

Abdominal aortic aneurysms (AAA) are defined as a progressive weakening and dilatation of the aortic wall in the abdomen. The natural course of the disease is to grow and rupture, which is associated with a mortality rate up to 90% [1]. AAA patients with a risk of rupture, currently determined solely based on aortic diameter, receive open or endovascular surgery. However, the predictive value of aortic diameter for rupture is limited, since rupture also occurs in patients with small aneurysms, while larger AAA can remain unruptured [2, 3]. Therefore, new indicators to better predict disease progression based on pathophysiologic mechanisms underlying AAA are warranted. Moreover, insight into AAA pathophysiology can contribute to novel non-invasive treatment options for aortic wall weakening.

Vascular smooth muscle cells (vSMC) are the predominant cell type within the aortic wall. These cells are radially oriented into multiple layers, embedded into the extracellular matrix (ECM). vSMC maintain functional and structural integrity of the aorta, and regulate blood flow and pulse pressure by using their contractile properties [4]. Force generation by vSMC is regulated via the actomyosin apparatus, consisting of thin filaments of polymerized smooth muscle α-actin (encoded by *ACTA2*), and myosin thick filaments, composed of a heavy chain dimer (encoded by *MYH11*), two essential and two regulatory light chains. Since mutations in genes encoding for these contractile proteins, including *ACTA2* and *MYH11,* cause hereditary thoracic AA (TAA) [5–7], impaired vSMC contractile capacity is also implied to have a role in sporadic AAA pathophysiology. Previous studies indeed show a reduction of known contractile vSMC markers, such as smooth muscle α-actin, calponin and smoothelin, in AAA tissue [8–10]. To further investigate the concept of vSMC contractile dysfunction in the AAA wall, measuring the actual contraction forces of vSMC derived from AAA patients is necessary.

We previously established a high throughput method to measure *in vitro* vSMC contractility using Electric Cell-substrate Impedance Sensing (ECIS) [11]. Our findings revealed impaired contraction in vSMC derived from AAA patients compared to healthy control vSMC [12]. Surprisingly, and contrary to our initial hypothesis, no correlation between vSMC contractility and the expression of known markers of the contractile vSMC phenotype was observed [12]. This indicates that *in vitro* vSMC contractility is not solely defined by the expression of known contractile markers or phenotypic state, and other yet unidentified proteins are implicated in impaired contraction of vSMC derived from AAA patients.

The aim of this study is therefore to uncover the underlying mechanism of altered contractility of vSMC derived from AAA patients and identify the proteins involved in this cascade. Quantification of vSMC contractile capacity using ECIS revealed large variation in contraction of vSMC derived from human AAA tissue (AAA-SMC) compared to vSMC derived from healthy aortic biopsies (C-SMC). This variability was used to define three AAA groups with low, intermediate (i.e. comparable to C-SMC contraction) and high contraction. Phosphoproteomic analysis on these cells identified NUAK family kinase 1 (NUAK1) as a novel regulator of contraction in AAA-SMC. Follow-up experiments were performed to further understand the mechanism and validate the data. Gaining more insight into the function of NUAK1 within the aortic wall contributes to further understanding of AAA pathophysiology.

## 2. Methods

### 2.1 Patient population and tissue collection

Biopsies at the largest diameter of the aorta were obtained from 24 AAA patients during open aneurysm repair in the Amsterdam University Medical Center, the Netherlands. All patients were above the age of 18 and signed informed consent to participate in the study. Control aortic biopsies were obtained from non-dilated infrarenal aortas of 8 *post-mortem* kidney donors without cardiovascular problems. Biopsies were transported directly after collection on ice-cold sterile 0.9% NaCl solution from the operating room to the laboratory. The tissue collection was fulfilled in accordance with the regulations of the WMA Declaration of Helsinki and institutional guidelines of the Medical Ethical Committee of the VU Medical Center (Biobank 2017.121: Aortic Aneurysms, Atherosclerosis and Biomarkers). Clinical characteristics of the study population used in the phosphoproteomic screen are reported in Table 1. As the kidney donors remained anonymous, only age and sex were reported for the control group.

**Table 1:** Clinical characteristics.

|  | C-SMC (n=8) | AAA-SMC Low (n=7) | AAA-SMC Intermediate (n=11) | AAA-SMC High (n=6) | P-value |
| --- | --- | --- | --- | --- | --- |
| **Age (years)** | 52.4 (16.7) | 71.7 (7.9) | 73.9 (9.7) | 71.8 (9.5) | **0.003** |
| **Sex (% male)** | 57.1 | 71.4 | 80 | 33.3 | 0.280 |
| **Aneurysm size (mm)** | N/A | 70.6 (16.8) | 73.8 (22.3) | 77.0 (17.0) | 0.838 |
| **Rupture (%)** | N/A | 14.3 | 10 | 33.3 | 0.475 |
| **Current smoking (%)** | NK | 42.9 | 11.1 | 33.3 | 0.341 |
| **Hypertension\* (%)** | NK | 71.4 | 33.3 | 80 | 0.155 |
| **Previous vascular surgery<sup>†</sup> (%)** | NK | 42.9 | 33.3 | 60 | 0.627 |
| **Renal dysfunction\* (%)** | N/A | 42.9 | 11.1 | 20 | 0.326 |
| **DM (%)** | NK | 28.6 | 10 | 33.3 | 0.478 |
| **BMI** | NK | 26.7 (5.7) | 28.2 (4.4) | 25.1 (4.0) | 0.463 |

Data are presented as *n* (%) or mean ± standard deviation. Groups were compared using one-way ANOVA or Kruskal-Wallis test when not normally distributed, with Bonferroni correction. Categorical data was compared with the Chi-square test. Abbreviations: C-SMC: Control smooth muscle cells, AAA-SMC: Abdominal aortic aneurysm smooth muscle cells, N/A = not applicable, NK = not known, DM = diabetes mellitus, BMI = body mass index
\*Hypertension and renal dysfunction were defined as diagnosed by a medical doctor or use of specific medication. The renal function of controls is presumed to be sufficient, as they have been approved for kidney donation.
<sup>†</sup>Previous vascular surgery includes percutaneous coronary intervention, percutaneous transluminal angioplasty, coronary artery bypass grafting and endovascular aneurysm repair.

### 2.2 Aortic biopsy explant and primary vSMC isolation

vSMC were isolated from aortic biopsies as described in Bogunovic *et al* [11]. In short, the biopsies were processed in a laminar flow cabinet, using sterilized surgical instruments. The adventitial layer and the intimal layer were removed from the biopsy, leaving a homogeneous and compact medial layer. This medial layer was sliced into approximately 7 to 8 pieces, which were placed on the bottom of a T-25 flask filled with 1.5 ml culture medium. vSMC were cultured in a humidified incubator at 37 °C and 5% CO₂, in Human Vascular Smooth Muscle Cell Basal Medium (Gibco, Thermo Fisher Scientific, Walthem, MA, USA) supplemented with Smooth Muscle Growth Supplement (Gibco, Thermo Fisher Scientific), 100 units/ml penicillin and 100 µg/ml streptomycin. Cell growth was observed over time around the tissue pieces and culture medium was refreshed once a week until 80-90% confluence was reached. The vSMC were then transferred into a T-75 flask using 0.05% trypsin-EDTA, and culturing was continued until confluence was reached again to use the cells for experiments. Primary vSMC were used between passages 1-8 in all experiments.

### 2.3 Cell contraction measurements

vSMC contraction measurements were performed as reported before using Electric Cell-Substrate Impedance Sensing (ECIS; Applied BioPhysics Inc, Troy, NY, USA) [11]. ECIS is a technique used to perform real-time contraction measurements of adherent cells *in vitro* by measuring impedance value based on electrode coverage. A sterile 96-well plate with 10 gold electrodes per well (96W10idf PET; Ibidi, Planegg, Germany) was prepared by coating the wells with 10 mM L-cysteine at room temperature (RT) for 30 min. After removing the L-cysteine, the plate was washed twice with sterile water. Subsequently, the plate was coated with 1% sterile gelatin solution per well and incubated for at least 1 h at 37 °C. vSMC were seeded in a density of 30,000 cells/well in 200 µl complete medium and the ECIS plate was installed in the ECIS 96-well plateholder in the incubator. Cells were allowed to attach for approximately 48 h, thereby forming a stable monolayer with a corresponding resistance value. The electrical impedance was recorded every 8 sec at a single frequency of 4 kHz.

To induce a contractile response, vSMC were treated with 10 µg/ml ionomycin (Sigma Aldrich, Darmstadt, Germany), which is a Ca²⁺ ionophore facilitating the intracellular transfer of calcium. Rapid vSMC contraction after stimulation is indicated by a drop in impedance value, caused by a reduction in electrode coverage by the cells. Cells were seeded in triplicate to account for intra-experimental variation in each individual experiment. Three independent experiments were performed to account for inter-experimental variation. Equation (1) was used to calculate the contractile response. C indicates the percentage of change in contraction compared to baseline, pre-stimulation (PrS) indicates the resistance value (expressed in Ohm) just upon ionomycin stimulation, and post-stimulation (PoS) indicates the resistance (expressed in Ohm) 2 min after finishing stimulation.

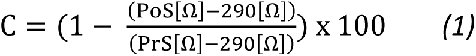

### 2.4 Phosphoproteomics analysis

#### 2.4.1 Sample collection

Each vSMC line was cultured in three 15-cm dishes until 70-80% confluency. After washing twice with cold PBS, cells were lysed (2 ml lysis buffer for 3 dishes) and detached using a cell scraper. Lysis buffer consisted of 20 mM HEPES pH 8.0, 9 M Urea (CH_4_N_2_O), 1 mM Orthovanadate (Na_3_VO_4_), 2.5 mM NaPP_i_ (Na_4_P_2_O_7_ (pyrophosphate)), 1 mM ß- Glycerophosphate (Na_2_C_3_H_7_PO_6_) in MilliQ. Lysates were sonicated on ice and debris was removed by centrifugation for 15 min at 5,400 x g at 17 °C. Protein concentration was measured using Pierce™ BCA Protein Assay Kit (Thermo Fisher Scientific, Waltham, MA, USA). For quality control, samples were loaded on a precast 4% to 12% NuPAGE Novex Bis-Tris 1.5 mm mini gel (Invitrogen). Electrophoresis was performed at 200 V in NuPAGE MES SDS running buffer until the dye reached the bottom of the gel. Subsequent, gels were fixed in 50% ethanol and 3% phosphoric acid solution and stained with 0.1% coomassie brilliant blue G-250 solution (34% methanol, 3% phosphoric acid, and 15% ammonium sulfate) (Supplementary Figure 1). Lysates were stored in -80 °C until the time of analysis.

#### 2.4.2 Sample preparation for phosphoproteomics

Lysates were thawed and insoluble material was removed by centrifugation. Samples were reduced with dithiotreitol (4 mM DTT, 30 min at 55°C) and alkylated with iodoacetamide (10 mM, 15 min in the dark). Next, the solution was diluted to 2 M urea by the addition of 20 mM HEPES pH 8.0 and digested with sequencing-grade modified trypsin (Promega) at a final concentration of 5 µg/ml overnight on RT. Digests were acidified with trifluoroacetic acid (TFA) to a final concentration of 0.1% and desalted using Oasis HLB cartridges (500 mg sorbent, Waters) after equilibration in 0.1% TFA. Bound peptides were washed twice with 0.1% TFA, eluted in 80% acetonitrile (ACN)/0.1% TFA, and lyophilized.

#### 2.4.3 IMAC enrichment and Nano-LC-MS/MS

Global phosphopeptide enrichment started from 200 µg peptides using IMAC cartridges (AssayMAP Fe(III)-NTA) on a Bravo Assaymap liquid handler (Agilent). Phosphopeptides were dried in a vacuum centrifuge and dissolved in 20 µl 0.5% TFA/4% ACN prior to injection; 18 µl was injected using partial loop injection. Peptides were separated using an Ultimate 3000 nanoLC-MS/MS system (Dionex LC-Packings, Amsterdam, The Netherlands) equipped with a 50-cm, 75-mm ID C18 Acclaim pepmap column (Thermo Scientific). After injection, peptides were trapped at 3 ml/min on a 10-mm, 75-mm ID Acclaim Pepmap trap column (Thermo Scientific) in buffer A (0.1% formic acid), and separated at 300 ml/min with a 10–40% buffer B (80% ACN/0.1% formic acid) gradient in 90 min (120 min inject-to-inject). Eluting peptides were ionized at +2 kV and introduced into a Q Exactive HF mass spectrometer (Thermo Fisher, Bremen, Germany). Intact masses were measured in the Orbitrap cell with a resolution of 120,000 (at m/z 200) using an automatic gain control (AGC) target value of 3 x 106 charges. The top 15 highest signal peptides (charge states 2+ and higher) were submitted to MS/MS in the higher energy collision cell (1.6-Da isolation width, 25% normalized collision energy). MS/MS spectra were measured in the Orbitrap with a resolution of 15k (at m/z 200) using an AGC target value of 1 x 106 charges and an underfill ratio of 0.1%. Dynamic exclusion was used with a repeat count of 1 and an exclusion time of 30 sec.

#### 2.4.4 Protein and phosphopeptide quantification

MS/MS spectra were searched against a Swissprot reference proteome (human, 2021_01 canonical plus isoforms, 42383 entries) using MaxQuant 1.6.10.43. Enzyme specificity was set to trypsin and up to two missed cleavages allowed. Cysteine carboxamidomethylation (+57.021464 Da) was treated as a fixed modification and serine, threonine and tyrosine phosphorylation (+79.966330 Da), methionine oxidation (+15.994915 Da) and N-terminal acetylation (+42.010565 Da) as variable modifications. The lysate MS/MS spectra were searched similarly, but without variable serine, threonine and tyrosine phosphorylation. Peptide precursor ions were searched with a maximum mass deviation of 4.5 ppm and fragment ions with a maximum mass deviation of 20 ppm. Peptide, protein and site identifications were filtered at an FDR of 1% using the decoy database strategy. The minimum peptide length was set at 7 amino acids, the minimum Andromeda score for modified peptides was 40, and the corresponding minimum delta score was 6 (default MaxQuant settings). Identifications were propagated across samples with the match between runs option checked.

#### 2.4.5 Phosphoproteomics data analysis

Protein-level differential analyses were performed using the label-free quantitation (LFQ) intensities in the proteinGroups.txt output table of the MaxQuant search. The counts of the proteomics data were normalized to the total spectral count per sample and the intensities of the global phosphoproteomics screen were normalized to the median intensity per sample. p<0.05 was considered significantly different. Significantly different data was further filtered to find the most discriminatory changes and to filter out low-level proteins that could not be reliably quantified. Proteomics data was filtered for proteins found in ≥50% of the samples in all four groups, and further selection was performed based on 1) Average normalized count >1.4, 2) Function of the protein and 3) Expression pattern correlated with vSMC contraction. Protein networks were generated using the STRING database (Search Tool for the Retrieval of Interacting Genes/Proteins) and visualized with Cytoscape software. Protein interaction networks were generated with ClusterONE and gene ontology (GO) analysis was performed using the BiNGO application in Cytoscape. Differential analyses at the phosphosite level were performed using the 1, 2, and 3 intensity levels in the Phospho (STY) Site.txt output table. MaxQuant search result tables were used to calculate Integrative iNferred Kinase Activity (INKA) scores, based on four parameters: the sum of all phosphorylated peptides belonging to a kinase; the detection of the phosphorylated kinase activation domain (kinase-centric parameters), the detection of known phosphorylated substrates and the presence of predicted phosphorylated substrates (substrate-centric parameters)[13]. The latest version of the INKA pipeline is available online at https://inkascore.org/.

### 2.5 Western Blot Analysis

Cells were washed with PBS and subsequently lysed in 100 µl SDS sample buffer (125 mM Tris-HCl pH 6.8, 4% SDS, 20% glycerol, 100 mM DTT, 0.02% Bromophenol Blue in MilliQ). After boiling the samples for 10 min at 95 °C, 15 µl of each sample was loaded. Proteins were separated by SDS-PAGE and subsequently transferred to nitrocellulose membranes. The membranes were blocked by incubating with BSA 5% for 2 h at RT on a rocking platform. Primary antibodies for Thrombospondin-1 (1:1000, #MA5-13398, Thermo Fisher Scientific), ATPase plasma membrane Ca^2+^ transporting 1 (1:1000, #PA1-914, Thermo Fisher Scientific), PDZ and LIM domain protein 4 (1:1000, NBP1-80833, Novus Biologicals), NUAK family kinase 1 (1:1000, #4458, Cell Signaling technology), phosphorylated Myosin light chain (Ser19, 1:1000, #3671, Cell Signaling Technology), phosphorylated Myosin phosphatase-targeting subunit 1 (Ser472, 1:1000, # PA5-114608, Thermo Fisher Scientific), Vinculin (1:1000, #V9131, Sigma-Aldrich), Myocardin (1:1000, SAB4200539, Sigma-Aldrich) and β-tubulin (1:2000, #2128, Cell Signaling Technology) or Glyceraldehyde-3-Phosphate Dehydrogenase (1:2000, #2118, Cell Signaling Technology) as loading control were incubated overnight at 4 °C. Secondary antibody incubation was performed for 1 h at RT with Polyclonal Goat Anti-Mouse Immunoglobulins/HRP (1:5000, Dako) or Polyclonal Goat Anti-Rabbit Immunoglobulins/HRP (1:5000, Dako), diluted in milk powder. Proteins were visualized with enhanced chemiluminescence (Amersham/GE Healthcare, Chicago, IL, USA) using the Amersham Imager 600 (GE Healthcare). Analysis of the band intensities was performed using ImageQuantTL.

### 2.6 RNA isolation and quantitative PCR

Gene expression levels were measured using quantitative PCR (qPCR). vSMC were cultured in a 12-well culture plate and lysed using 300 µl RNA lysis buffer (Zymo Research, Irvine, CA, USA). RNA was isolated with the Quick-RNA Miniprep Kit (Zymo Research) and synthesis of complementary DNA (cDNA) was performed in a reverse transcription reaction using SuperScript VILO cDNA Synthesis Kit (Thermo Fisher Scientific). To quantify gene expression, qPCR was performed using the CFX384 Touch Real-Time PCR System (Bio-Rad Laboratories Inc, CA, USA) with the use of LightCycler SYBR Green I Master (Roche Applied Science, Penzberg, Germany). The 2-deltaCT method was used to calculate relative gene expression levels between the different samples. TATA-Box Binding Protein (*TBP*) was used as housekeeping gene. Primer sequences are provided in Supplementary Table 1.5.

### 2.7 siRNA transfection

Cells were cultured until 70 to 80% confluency. siRNA transfection using Lipofectamine™ 2000 Transfection Reagent (Thermo Fisher Scientific) in OptiMEM (Gibco) was performed. For gene silencing, a final concentration of 150 nM of siRNA (Thrombospondin-1 siRNA ID: s14099, PDZ and LIM domain protein 4 siRNA ID: s16326, ATPase plasma membrane Ca^2+^ transporting 1 siRNA ID: s752, NUAK family kinase 1 siRNA ID: s90, Thermo Fisher Scientific) was used. Silencer™ Select Negative Control No. 1 siRNA (Thermo Fisher Scientific) was used as negative control. Cells were used for experiments 24 h after transfection.

### 2.8 Immunofluorescence staining and microscopy

Cells were cultured on sterile glass coverslips (Ø 13 mm, Thermo Fisher Scientific). After rinsing with EBSS, cells were fixated with 4% formaldehyde for 10 minutes at RT. Next, coverslips were washed three times with PBS 0,05% Tween (PBST) and cell membranes were permeabilized with 0.2% Triton X-100 in PBS for 10 minutes. The samples were incubated with blocking solution, consisting of 1% human serum albumin (HSA) in PBST for 1 h at RT. Cells were incubated with DAPI (1:200, #62248, Thermo Fisher Scientific) and Actin-stain 670 Phalloidin (1:100, #PHDN1-A, Cytoskeleton) for 1 h at RT. Slides were closed with Mowiol4- 88/DABCO solution (Calbiochem, Sigma Aldrich). Images were acquired using the Nikon A1R (Nikon, Tokyo, Japan) microscope and the corresponding software Nis-Elements C Sofware (Nikon). Representative images were analyzed and adjusted using ImageJ.

### 2.9 Statistical Analysis

All analyses were performed with the use of SPSS version 28.0.1.1 (IBM Statistics, Armonk, NY, USA). Two groups were compared using (paired) T-test and multiple groups were compared using one-way ANOVA. When data was not normally distributed, non-parametric tests were used (two groups: Mann-Whitney U test, multiple groups: Kruskal-Wallis test, Bonferroni correction). Correlations were analyzed with the Pearson correlation or Spearman’s Rank correlations when not normally distributed. Categorical data was compared with the Chi-square test. Data are presented with mean and standard deviation indicated by the bars. A value of p≤0.05 was considered statistically significant. Graphs were constructed with the use of GraphPad Prism 9.1.0 (GraphPad Software, La Jolla, CA, USA).

## 3. Results

### 3.1 Large variability in AAA-SMC contraction compared to C-SMC contraction

Using our new method to measure vSMC contractility, we observed large variability in contraction of AAA-SMC compared to contraction of C-SMC (Figure 1A). Therefore, the mean contractility of C-SMC ±2 standard deviations (SD) was calculated as a reference for normal vSMC contractility (83,3% ± 6,3%). Based on this C-SMC range, three AAA-SMC groups were defined with low, intermediate (i.e. comparable to C-SMC) and high contraction (Figure 1B, AAA-SMC Low contracting: mean±SD: 70,1±5,7% (n=7); AAA-SMC Intermediate contracting: 83,5±2,7% (n=11); AAA-SMC High contracting: 92,0±1,3% (n=6)). In addition to the C-SMC and AAA-SMC used for the phosphoproteomics screen, the subdivision of the three AAA-SMC

**Figure 1:**
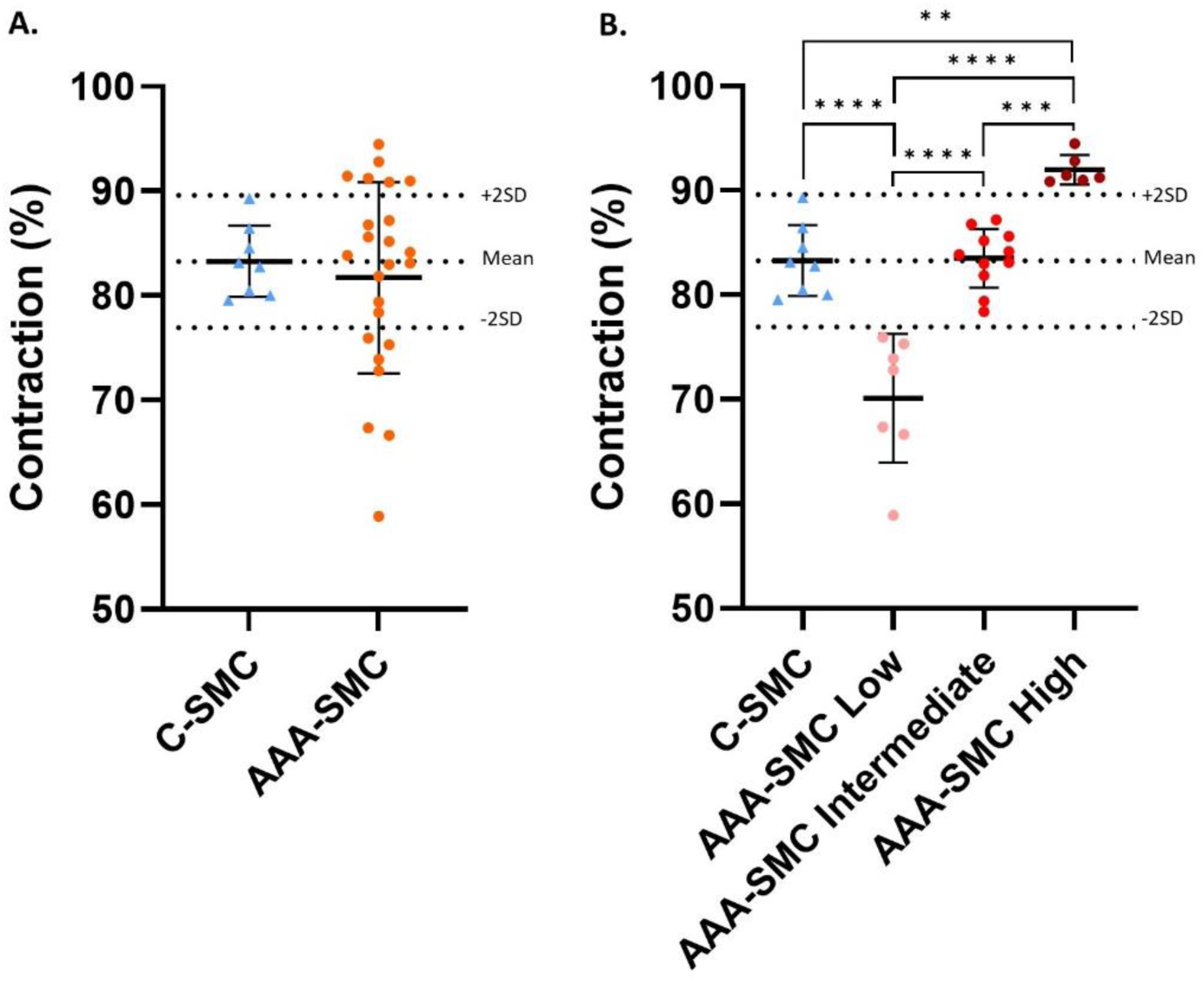
Contraction of C-SMC and AAA-SMC. A. Contraction of C-SMC (n=8) and AAA-SMC (n=24) were measured using Electric Cell-substrate Impedance Sensing. B. AAA-SMC were subdivided in AAA-SMC Low contracting (n=7), AAA-SMC Intermediate contracting (n=11) and AAA-SMC High contracting (n=6), based on the mean contractile response of C-SMC ±2SD, as represented by the dotted lines. Cells were seeded in triplicate within each individual experiment and three independent experiments were performed for each vSMC line. Two groups were compared using T-test and multiple groups were compared using one-way ANOVA, with Bonferroni correction. Data is presented as mean with error bars indicating standard deviations. Abbreviations: C-SMC: Control smooth muscle cells, AAA-SMC: Abdominal aortic aneurysm smooth muscle cells, SD: Standard deviation, **p<0.01, ***p<0.001, ****p<0.0001.

groups based on contractile response was verified in a larger cohort (in total; C-SMC n=18 and AAA-SMC n=39, Supplementary Figure 2).

### 3.2 Clinical characteristics of the C-SMC and three AAA-SMC groups

Table 1 shows the clinical characteristics of the C-SMC and three AAA-SMC groups. Age at the time of biopsy was significantly lower in C-SMC compared to all three AAA-SMC groups, but did not correspond to vSMC contraction (Supplementary Figure 3). Sex was equally distributed between the four groups. Other clinical characteristics, including aneurysm size, rupture, smoking status, hypertension, previous vascular surgery, renal dysfunction, diabetes mellitus and BMI, were similar in the three AAA-SMC groups.

### 3.3 Proteomic analysis shows that abundance of Thrombospondin-1, ATPase plasma membrane Ca^2+^ transporting 1 and PDZ and LIM domain protein 4 correlated with vSMC contraction

Hierarchical clustering of the proteomics data revealed that the majority of samples from the AAA-SMC Intermediate contracting group were most similar to the C-SMC group, which corresponds to their contractile ability (Figure 2A). In total, 4041 proteins were identified in the proteomics screen (Supplementary Table 1.1). A Venn diagram demonstrates that 3129 proteins were present in all four groups, as well as unique proteins found in each group (Figure 2B). When comparing the four groups, 60 proteins were found to differ significantly in their expression level (Supplementary Table 1.2). Figure 2C shows the protein interaction network of these 60 proteins and the separate clusters associated with the following GO terms are shown in Supplementary Figure 4: Cell-substrate adhesion (GO ID: 31589, 8 nodes), Deoxyribose phosphate metabolic process (GO ID: 19692, 4 nodes), Cell adhesion (GO ID: 7155, 5 nodes) and Negative regulation of fibroblast apoptotic processes (GO ID: 2000270, 5 nodes).

**Figure 2:**
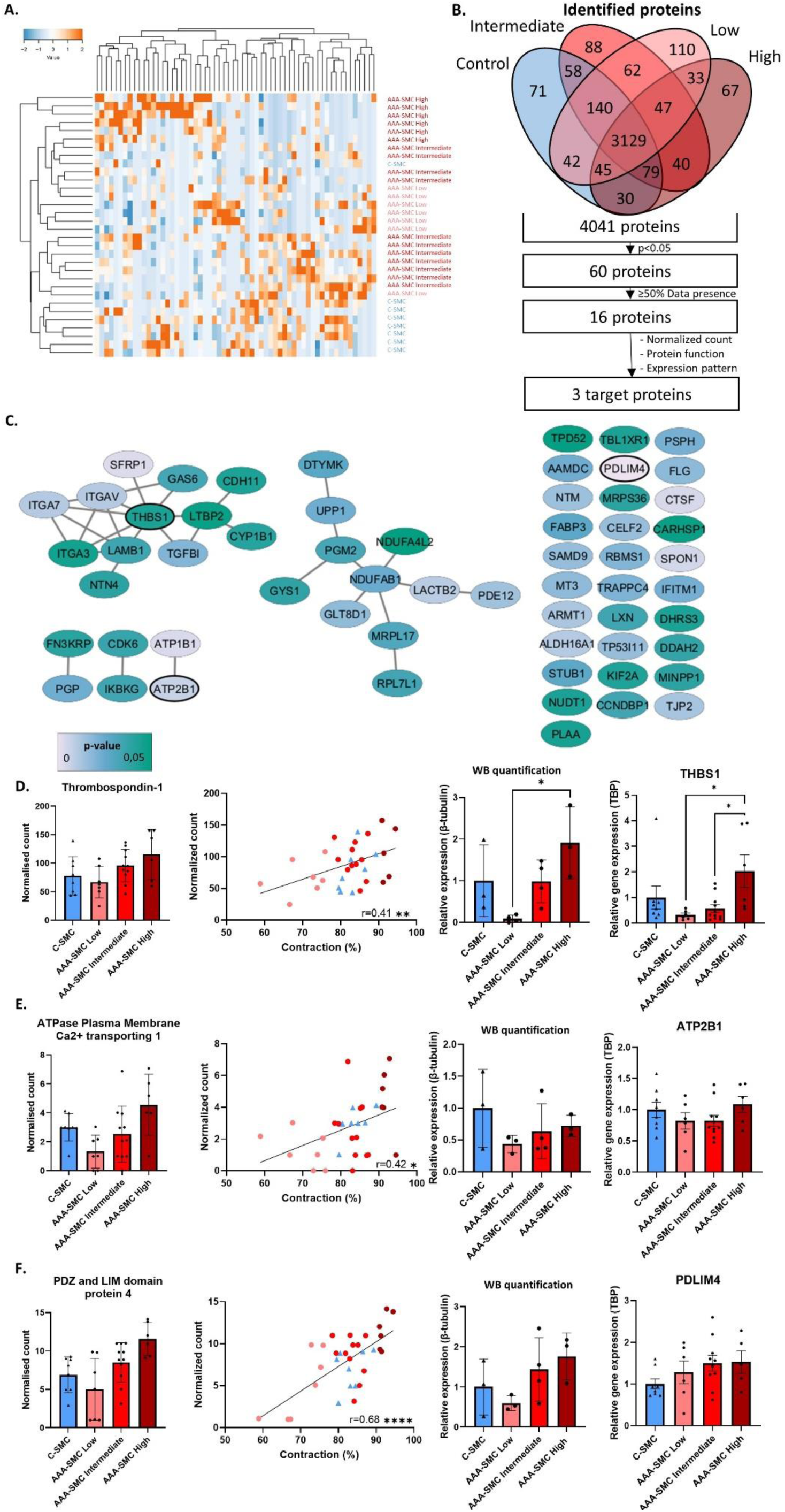
Proteomic analysis shows that the expression of Thrombospondin-1, ATPase plasma membrane Ca^2+^ transporting 1 and PDZ and LIM domain protein 4 correlated with vSMC contraction. A. Hierarchical clustering of all proteins that are differentially expressed at p<0.05 between the four groups (C-SMC n=8, AAA-SMC Low contracting n=7, AAA-SMC Intermediate contracting n=11 and AAA-SMC High contracting n=6). B. Venn diagram showing the number of identified proteins within each of the four groups. 60 proteins were significantly differentially expressed between the four groups and these were further filtered based on ≥50% data presence within each group, resulting in 16 proteins. Proteins were filtered for their normalized count, protein function and expression pattern, resulting in three target proteins. C. Clustering of the 60 significantly different proteins. The three target proteins are shown with a thick outline. Colour of the nodes represent the p-value, as indicated in the figure. D. Normalized count of Thrombospondin-1 as measured in the proteomics screen, correlation of the Thrombospondin-1 count with contraction, bar graph showing quantification of Thrombospondin-1 expression on WB to verify the proteomics data, and bar graph showing THBS1 gene expression measured using qPCR in the same C-SMC and AAA-SMC as used in the proteomics screen. E. Normalized count of ATPase Plasma Membrane Ca^2+^ transporting 1 as measured in the proteomics screen, correlation of the ATPase Plasma Membrane Ca^2+^ transporting 1 count with contraction, bar graph showing quantification of ATPase Plasma Membrane Ca^2+^ transporting 1 expression on WB to verify the proteomics data, and bar graph showing ATP2B1 gene expression measured using qPCR in the same C-SMC and AAA-SMC as used in the proteomics screen. F. Normalized count of PDZ and LIM domain protein 4 as measured in the proteomics screen, correlation of the PDZ and LIM domain protein 4 count with contraction, bar graph showing quantification of PDZ and LIM domain protein 4 expression on WB to verify the proteomics data, and bar graph showing PDLIM4 gene expression measured using qPCR in the same C-SMC and AAA-SMC as used in the proteomics screen. Full WB pictures of quantified proteins in 2D-F are available in Supplementary Figure 5. Groups were compared using one-way ANOVA or Kruskal-Wallis test when not normally distributed, with Bonferroni correction. Correlations were analyzed with the Pearson correlation or Spearman’s Rank correlations when not normally distributed. Data is presented as mean with error bars indicating standard deviations. Abbreviations: C-SMC: Control smooth muscle cells, AAA-SMC: Abdominal aortic aneurysm smooth muscle cells, WB: Western blot, qPCR: quantitative PCR, TBP: TATA-Box Binding Protein, *p<0.05, **p<0.01, ****p<0.0001.

To focus on differential proteins with widespread expression, an ≥50% data presence filter was applied, resulting in 16 proteins (Figure 2B). These 16 proteins were further selected based on 1) Average normalized count >1.4, 2) Function of the protein and 3) Expression pattern correlated with vSMC contraction. This resulted in three target proteins, namely Thrombospondin-1 (encoded by *THBS1*), ATPase plasma membrane Ca^2+^ transporting 1 (encoded by *ATP2B1*) and PDZ and LIM domain protein 4 (encoded by *PDLIM4*) (Figure 2D-F).

The expression of these three proteins correlated with vSMC contraction, with low expression in the low contracting AAA-SMC and high expression in the high contracting AAA-SMC (Figure 2D-F). Alterations in protein expression levels were verified on western blot (Quantification in Figure 2D-F, Western blots in Supplementary Figure 5). Gene expression levels of *THBS1, PDLIM4* and *ATP2B1* were measured using qPCR and aligned with protein expression levels (Figure 2D-F).

To investigate a mechanistic role of these three proteins in vSMC contraction, siRNA-mediated knockdown (KD) was performed, after which vSMC contraction was measured again using ECIS. Although efficiency of the siRNA-mediated KD was verified (Supplementary Figure 6), no reproducible reduction in vSMC contraction was found after KD.

### 3.4 Phosphoproteome of AAA-SMC reveals NUAK1 as a potential regulator of contraction

A Venn diagram demonstrates that 10713 phosphosites were identified as in common for all four groups, as well as unique phosphosites in each group (Figure 3A). Of the 16997 identified phosphosites (Supplementary Table 1.3), 396 phosphosites were found to be differentially phosphorylated when comparing the four groups (Figure 3A and Supplementary Table 1.4). Cluster and GO analysis on the proteins belonging to the phosphopeptides was performed to get insight into processes potentially involved in disturbed contraction by changes in phosphorylation levels. The three largest clusters were associated with the following GO terms: Integrin activation (GO ID: 33622, 10 nodes), Ribosome biogenesis (GO ID: 42254, 9 nodes) and Actin cytoskeleton organization (GO ID: 30036, 8 nodes) (Clusters in Supplementary Figure 7).

**Figure 3:**
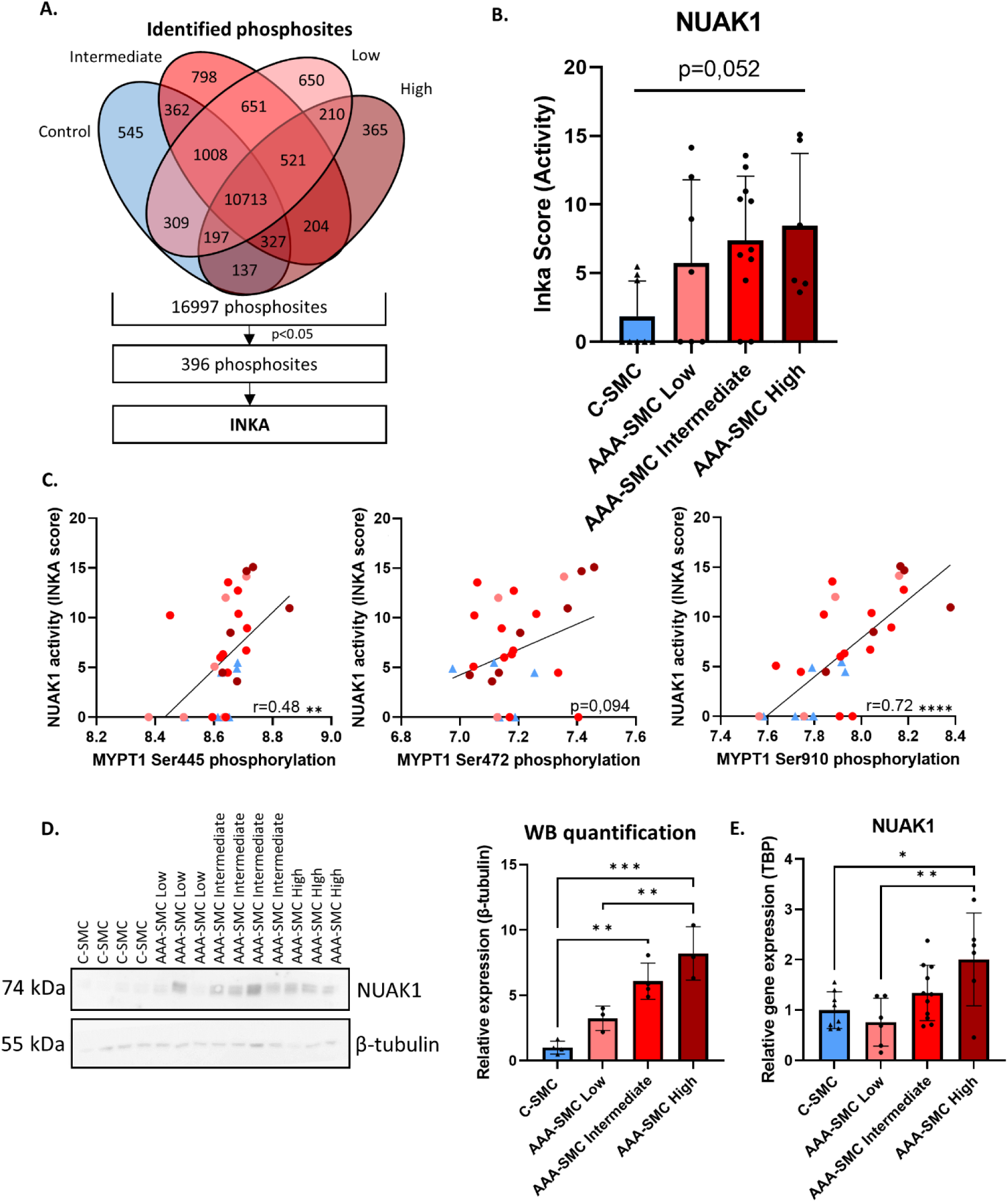
Phosphoproteome of AAA-SMC reveals NUAK1 as potential regulator of contraction. A. Venn diagram showing the number of identified phosphosites within each of the four groups. 394 phosphosites were significantly differently phosphorylated between the four groups. INKA analysis was performed on this dataset to find differences in kinase activity between the four groups. B. INKA scores of NUAK1 in the four groups (C-SMC n=8, AAA-SMC Low contracting n=7, AAA-SMC Intermediate contracting n=11 and AAA-SMC High contracting n=6). C. Correlation between phosphorylation of three MYPT1 phosphosites (Ser445, Ser472, Ser910) and NUAK1 activity. D. WB picture and quantification of NUAK1 protein levels in C- SMC (n=4), AAA-SMC Low contracting (n=3), AAA-SMC Intermediate contracting (n=4) and AAA-SMC High contracting (n=3). E. RNA expression levels of NUAK1 quantified with qPCR in C-SMC (n=8), AAA-SMC Low contracting (n=7), AAA-SMC Intermediate contracting (n=11) and AAA-SMC High contracting (n=6). Groups were compared using one-way ANOVA or Kruskal- Wallis test when not normally distributed, with Bonferroni correction. Correlations were analyzed with the Pearson correlation or Spearman’s Rank correlations when not normally distributed. Data is presented as mean with error bars indicating standard deviations. Abbreviations: C-SMC: Control smooth muscle cells, AAA-SMC: Abdominal aortic aneurysm smooth muscle cells, NUAK1: Nu (novel) AMPK related kinase, INKA: Integrative Inferred Kinase Activity, MYPT1: Myosin phosphatase targeting subunit 1, qPCR: quantitative PCR, WB: Western Blot, TBP: TATA-Box Binding Protein, *p<0.05, **p<0.01, ***p<0.001, ****p<0.0001.

Based on the identified phosphosites, INKA analysis was performed to find differences in kinase activity between the four group. Kinase activity of A-Raf Proto-Oncogene, Serine/Threonine Kinase (ARAF) was found to differ significantly (Supplementary Figure 8) and NUAK family kinase 1 (NUAK1) showed a trend towards significantly increased kinase activity in higher contracting AAA-SMC (Figure 3B). Since NUAK1 is known to modulate activity of Myosin phosphatase-targeting subunit 1 (MYPT1), a protein involved in regulation of the actomyosin apparatus, this kinase was of specific interest. Phosphorylation on three MYPT1 amino acids (Ser445, Ser472 and Ser910) by NUAK1 inhibits MYPT1 phosphatase activity [14]. Subsequently, dephosphorylation of Myosin light chain (MLC) by MYPT1 is prevented and this regulates contractile function in various cell types. This role of NUAK1 to phosphorylate MYPT1 was confirmed in our data, since a correlation was found between NUAK1 activity and phosphorylation levels of two MYPT1 phosphosites (Figure 3C). The phosphorylation levels of the third phosphosite (Ser472) showed a positive trend towards correlation with NUAK1 activity (Figure 3C). Looking further downstream, phosphorylation levels of MLC (pMLC, Ser19 and Thr18+Ser19) found in the phosphoproteomics data were not correlated with vSMC contraction; the same was observed on western blot (Supplementary Figure 9). NUAK1 protein expression was not detected within the proteomics screen, but protein levels measured using western blot and RNA expression both showed a similar expression pattern as NUAK1 activity (Figure 3D-E).

### 3.5 NUAK1 regulates contraction in AAA-SMC

To validate the role of NUAK1 activity in vSMC contraction, experiments were performed in a second, independent cohort of C-SMC and AAA-SMC. NUAK1 KD (KD validation in Figure 4B and Supplementary Figure 6 and 12) decreased contraction in AAA-SMC (Figure 4A). Contraction of C-SMC remained within the ±2SD range after NUAK1 KD (Figure 4A), indicating a unique NUAK1 dependent contraction mechanism in AAA-SMC. Since NUAK1 can phosphorylate MYPT1, which subsequently might affect pMLC, changes in pMYPT (Ser472) and pMLC (Ser19) after NUAK1 KD were examined. A decrease of pMLC expression after NUAK1 KD was seen in C-SMC and AAA-SMC (Figure 4B). pMYPT levels were reduced in the majority of C-SMC and AAA-SMC after NUAK1 KD, but on average there was no significant reduction (Figure 4B).

**Figure 4:**
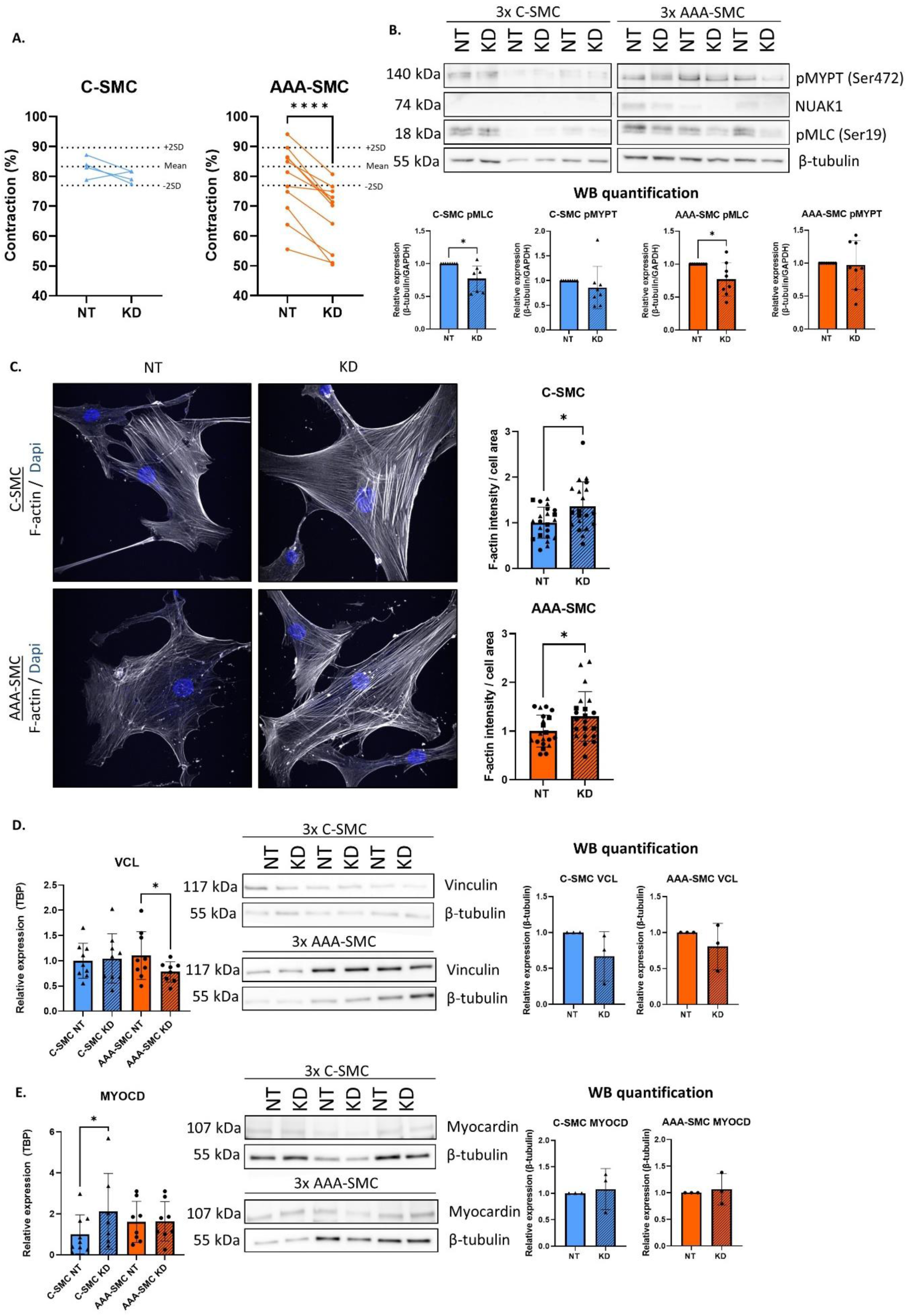
NUAK1 regulates contraction in AAA-SMC. A. Contraction of C-SMC (n=4) and AAA- SMC (n=10) after NUAK1 KD for 24 hours measured using electric cell substrate impedance sensing. B. Effect of NUAK1 KD on pMYPT (Ser472) and pMLC (Ser19) shown in representative WB picture and quantified in C-SMC (n=8) and AAA-SMC (n=8). C. Representative immunofluorescence images showing F-actin (white) after NUAK1 KD in C-SMC and AAA-SMC. Nuclei/Dapi is shown in blue. Quantification of all images is shown in the graphs (each point represents an image and the different symbols indicate independent experiments in C-SMC (n=3) or AAA-SMC (n=3)). D. Effect of NUAK1 KD on VCL gene expression in C-SMC (n=12) and AAA-SMC (n=12), measured using qPCR, and protein expression in C-SMC (n=3) and AAA-SMC (n=3), as shown in WB picture and the quantification. E. Effect of NUAK1 KD on MYOCD gene expression in C-SMC (n=12) and AAA-SMC (n=12), measured using qPCR, and protein expression in C-SMC (n=3) and AAA-SMC (n=3), as shown in WB picture and the quantification. Two groups were compared using paired T-test. Data is presented as mean with error bars indicating standard deviations. Abbreviations: C-SMC: Control smooth muscle cells, AAA-SMC: Abdominal aortic aneurysm smooth muscle cells, NUAK1: Nu (novel) AMPK related kinase, KD: Knock down, NT: Non targeting siRNA, 2SD: ±2 standard deviations, pMYPT1: Phosphorylation of Myosin phosphatase targeting subunit 1, pMLC: Phosphorylation of Myosin light chain, VCL:Vinculin, MYOCD: Myocardin, GAPDH: Glyceraldehyde-3-Phosphate Dehydrogenase, TBP: TATA-Box Binding Protein, qPCR: quantitative PCR, WB: Western Blot, *p<0.05, ****p<0.0001.

Next to phosphorylation levels of MYPT1 and MLC, we further explored the underlying mechanism of altered vSMC contraction regulated by NUAK1. More stress fiber formation was observed after NUAK1 KD in both C-SMC and AAA-SMC, indicated by F-actin staining intensity (Figure 4C). Furthermore, *VCL* gene expression, encoding for the focal adhesion protein Vinculin, reduced in AAA-SMC after NUAK1 KD (Figure 4D). On protein level, a trend towards Vinculin reduction was seen after NUAK1 KD in C-SMC and AAA-SMC (Figure 4D). Myocardin (encoded by *MYOCD*) is a transcriptional activator inducing expression of vSMC contractile markers, such as smooth muscle α-actin, calponin and smoothelin [15]*. MYOCD* gene expression was increased after NUAK1 KD in C-SMC, which was not seen in AAA-SMC (Figure 4E). On protein level, increased Myocardin was seen in the majority of C-SMC after NUAK1 KD (Figure 4E). The expression of the contractile genes induced by Myocardin remained unchanged after 24 hours of NUAK1 KD (time point used in all experiments, Supplementary Figure 10). After 72 hours of NUAK1 KD, myosin light chain kinase 1 (encoded by *MYLK1*) gene expression increased and an increased trend in gene expression of *SMTN* and *CNN1* was measured (Supplementary Figure 10). Myocardin expression was not identified in the proteomics screen, but *MYOCD* gene expression showed a similar expression pattern as NUAK1 expression (Supplementary Figure 11).

The expression levels of multiple other genes, encoding for proteins which were previously described to be involved in vSMC contraction or related to NUAK1, were measured but did not change after NUAK1 KD in C-SMC and AAA-SMC. These genes included the three proteomics targets (*THBS1, PDLIM4, ATP2B1*), vSMC markers and contractile proteins (*ACTA2, MYLK1, CNN1, PPP1R12A, PRKG1, SMTN*), ECM organization proteins (*FN1, FBN-1, TNS1, SERPINE1, MMP2*), actin cytoskeleton (*FLNA, TLN1*), focal adhesion (*PXN*), collagens (*COL1A1, COL1A2, COL3A1*), integrins (*ITGA4, ITGA6, ITGB1, ITGB2, ITGAB3*), proliferation (*KI67*) and miscellaneous (*NUAK2, CDH11*) (Supplementary Figure 12).

## 4. Discussion

This study presents NUAK1 as a novel regulator of *in vitro* contraction in vSMC derived from AAA patients. We defined three AAA-SMC subgroups with low, intermediate and high contraction and phosphoproteomic analysis revealed that the contractile ability positively correlated with NUAK1 expression and activity. Mechanistically, NUAK1 KD reduced pMLC levels and *VCL* gene expression, and increased F-actin cytoskeletal filaments, along with decreased contraction in AAA-SMC (See overview Figure). Notably, NUAK1 KD did not impact contraction in C-SMC, suggesting that this novel mechanism is specific for vSMC derived from AAA patients.

NUAK1 belongs to a family of twelve AMPK-related kinases with great sequence homology to the catalytic domain of AMPK, all activated by the master upstream kinase Liver Kinase B1 (LKB1) [16, 17]. So far, this kinase has mainly been described in cancer research [18]. Our findings showed that NUAK1 expression and activity levels regulated contraction in AAA-SMC.

NUAK1 KD decreased contraction in AAA-SMC, combined with reduced pMLC levels. These lower pMLC levels are potentially caused by increased MYPT1 phosphatase activity due to decreased phosphorylation on MYPT1 Ser445, Ser472 and Ser910, as a result of low NUAK1 expression (See overview Figure). Interestingly, although the majority of AAA-SMC (and C- SMC) exhibited decreased MYPT1 Ser472 phosphorylation after NUAK1 KD, no overall significant decrease of MYPT1 Ser472 phosphorylation was observed after the KD. This discrepancy might be due to inter-individual variation among vSMC or the lack of significant correlation between NUAK1 activity and phosphorylation of MYPT1 on Ser472 specifically (Figure 3C). In line with our findings, NUAK1 was reported to regulate MYPT1 activity in HEK293 cells via phosphorylation on the three MYPT1 phosphosites, and subsequently, reduce dephosphorylation of MLC [14]. While our study represents the first evidence that NUAK1 regulates contraction in AAA-SMC, NUAK1 was found to have a similar role in human prostate stromal cells [19] and endothelial cells [20].

In addition to phosphorylation levels of MYPT1 and MLC, we aimed to identify other proteins regulated by NUAK1 and implicated in vSMC contraction. An increase in F-actin stress filaments was quantified after NUAK1 KD in both C-SMC and AAA-SMC. Since actin polymerization within vSMC is separately regulated from the actomyosin crossbridge cycling process [21], this increase in F-actin seen after NUAK1 KD might be an adaptive effect, consequent to changes in pMLC, making the cells more rigid. Consistent with our findings, knock out (KO) of NUAK1 in mouse embryonic fibroblasts (MEFs) increased F-actin filaments compared to wild type (WT) MEFs [14]. However, another study using the same cell type reported the opposite effect, namely decreased F-actin in LKB1 (master regulator of NUAK1) KO MEFs compared to WT MEFs, combined with decreased contraction (measured by cell wrinkling) [22]. Actin filaments within vSMC are linked to the ECM through focal adhesions and integrins, making the vSMC able to respond to changes in the ECM [23]. Reduced *VCL* gene expression, encoding for the focal adhesion protein Vinculin, was seen after NUAK1 KD in AAA-SMC. Vinculin reduction can impair the link between actin filaments and the ECM, which might induce the reported changes in actin cytoskeleton dynamics after NUAK1 KD, and subsequently affect mechanotransduction and contraction in AAA-SMC. The two aforementioned studies also reported altered Vinculin expression in LKB1/NUAK1 KO MEFs compared to WT MEFs [14, 22]. Taken together, these findings indicate that NUAK1 is involved in the regulation of actin cytoskeletal dynamics, via alterations in the focal adhesion protein Vinculin (as also implied by the GO terms “integrin activation” and “actin cytoskeleton organization” linked to the phosphoproteomics data (Supplementary Figure 7)), thereby affecting vSMC contractile capacity.

Noteworthy, high NUAK1 gene expression was also found in a TAA-SMC subgroup, identified using nuclear RNA sequencing on human TAA tissue compared to control tissue [24]. This TAA- SMC subgroup had the highest increase in proportion in TAA compared with control biopsies, and was characterized with the GO terms “muscle contraction” and “actin filament-based process”, compared to the other two smaller TAA-SMC subgroups associated with “ECM organization”. Moreover, NUAK1 was reported to regulate additional cellular processes known to contribute to aortic wall weakening, including regulation of matrix metalloproteinase (MMP)-2 and MMP-9 expression [25, 26], cell senescence and proliferation [27–29], fibrosis [30] and energy metabolism [31]. These previous findings combined with our study suggest that NUAK1 emerges as a promising target for further AAA research.

Despite the observed decrease in pMLC and increase in F-actin stress fibers, contraction remained unaffected after NUAK1 KD in C-SMC. This indicates a disease-related mechanism, in which contraction is NUAK1 dependent in AAA-SMC only. The pre-existing low NUAK1 activity and expression in C-SMC compared to AAA-SMC (Figure 3B and 3D-E) might explain why NUAK1 KD has no effect. Furthermore, other kinases can phosphorylate MYPT1 and MLC independently from NUAK1 [14, 16, 17, 32], as also shown by the detection of basal phosphorylation levels of MYPT1 and MLC. Additionally, Myocardin increased after NUAK1 KD in C-SMC and *MYOCD* gene expression exhibited a similar expression pattern as NUAK1 gene/protein expression. These observations together suggest that increased Myocardin might represent a compensatory mechanism aiming to restore the expression of NUAK1 and other contractile markers [15], but further experiments are warranted to verify this link between Myocardin and NUAK1.

The expression of Thrombospondin-1, ATPase plasma membrane Ca^2+^ transporting 1 and PDZ and LIM domain protein 4 correlated with vSMC contraction, but KD of these targets individually did not affect vSMC contraction. Nonetheless, since expression differences were found between AAA-SMC and C-SMC, we believe that these proteins are involved in AAA pathophysiology, as novel parts of known processes or as parts of yet unidentified pathological mechanisms. The adhesive glycoprotein Thrombospondin-1 mediates cell-to-cell and cell-to-matrix interactions and was described before to have a protective, as well as a promoting role in AAA development in human and rodent studies [33–39]. Such conflicting roles of Thrombospondin-1 may explain the varied levels observed in AAA-SMC in our study, and suggest that this protein may contribute more to AAA pathophysiology in a subset of patients. Additionally, Thrombospondin-1 has previously been linked to NUAK1; NUAK1 KO in epithelial ovarian cancer cells decreased *THBS1* gene expression [40]. While NUAK1 and Thrombospondin-1 demonstrated a similar expression pattern in our data, NUAK1 KD did not affect *THBS1* levels. Furthermore, PDZ and LIM domain protein 4, involved in actin cytoskeleton reorganization and cell migration, has not been described in the context of AAA before. However, its homolog PDLIM5 was found to suppresses the expression of contractile proteins in pulmonary arterial SMC [41]. The third protein, ATPase plasma membrane Ca^2+^ transporting 1, catalyzes the hydrolysis of ATP coupled with the transport of calcium from the cytoplasm to the extracellular space, thereby maintaining intracellular calcium homeostasis, which is crucial for vSMC contraction [42].

## Strengths and limitations

AAA have a broad spectrum of potential causes, including lifestyle, sex or other cardiovascular disorders, and our samples therefore provide a broad and yet realistic, and thus representative insight into the phosphoproteome of vSMC derived from an AAA patients. Despite heterogeneity and variation between patients, our study identified NUAK1 as a novel regulator of contraction of vSMC derived from all AAA patients used in this study. Focus on additional cellular processes regulated by NUAK1 within the aortic wall in future research can provide a more comprehensive understanding of the disease mechanism. Limitations include the overall younger age and lack of other clinical characteristics within the control group compared to the AAA group. However, no correlation between age and vSMC contraction was observed (Supplementary Figure 2). Additionally, the use of an explant protocol to obtain vSMC and culturing cells *in vitro* in a new environment without flow or pressure might influence their properties or phenotype and may not fully reflect the *in vivo* situation.

## Conclusion

NUAK1 expression and activity levels correspond to the contractile ability of AAA-SMC. Low NUAK1 increased MYPT1 phosphatase activity, and subsequent lower pMLC levels coincided with reduced contraction in AAA-SMC. Impaired vSMC contraction within the aortic wall might decrease the ability to resist blood flow and pressure, leading to aortic wall weakening and subsequent AAA development. Further *in vitro* and *in vivo* studies are necessary to gain a complete understanding of the mechanisms and function of NUAK1 within the aortic wall, in both normal and pathological context.

## Funding

This work was supported by the Dutch Heart Foundation/Hartstichting, Dekkerbeurs 2019T065 Senior Clinical Scientist grant.

### Author Contribution Statement

KBR designed and planned the study, performed the experiments, analyzed the data and wrote the manuscript. AAH, JCK, TVP, SRP, CRJ advised and contributed to the design, experiments and analysis of the phosphoproteomics data set. TARM, PLH, JvdV, NB, KKY helped with the research design, advised on the experiments and revised the manuscript. All authors reviewed the manuscript.

## Acknowledgments

We gratefully acknowledge the surgical team from the Amsterdam UMC for providing the aortic material during surgery and Lan Tran and Samira van Knippenberg for biobanking support. We wish to thank Albert van Wijk for experimental support. Netherlands Organisation for Scientific Research (NWO-Middelgroot project number 91116017 to CRJ) is acknowledged for support of the mass spectrometry infrastructure and Surfsara for computing infrastructure (reference e-infra180166).

