## Supplemental Figure 1-12 for "NUAK1 regulates contraction in vascular smooth muscle cells derived from abdominal aortic aneurysm patients"

### Supplementary Figures

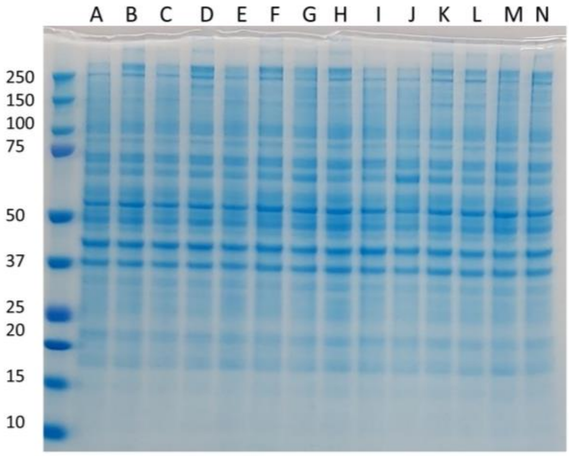


| Slot | Letter | Group |
|---|---|---|
| **1** | **Ladder** | |
| **2** | A | AAA-SMC Low |
| **3** | B | AAA-SMC Intermediate |
| **4** | C | AAA-SMC High |
| **5** | D | C-SMC |
| **6** | E | AAA-SMC Low |
| **7** | F | AAA-SMC Intermediate |
| **8** | G | AAA-SMC High |
| **9** | H | C-SMC |
| **10** | I | AAA-SMC Low |
| **11** | J | AAA-SMC Intermediate |
| **12** | K | AAA-SMC High |
| **13** | L | C-SMC |
| **14** | M | AAA-SMC Low |
| **15** | N | AAA-SMC Intermediate |

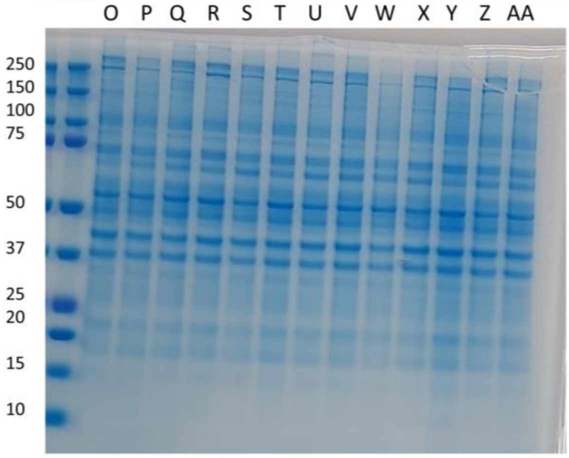


| Slot | Letter | Group |
|---|---|---|
| **1** | **Ladder** | |
| **2** | **Ladder** | |
| **3** | O | AAA-SMC High |
| **4** | P | C-SMC |
| **5** | Q | AAA-SMC Low |
| **6** | R | AAA-SMC Intermediate |
| **7** | S | AAA-SMC High |
| **8** | T | C-SMC |
| **9** | U | AAA-SMC Low |
| **10** | V | AAA-SMC Intermediate |
| **11** | W | AAA-SMC High |
| **12** | X | C-SMC |
| **13** | Y | AAA-SMC Low |
| **14** | Z | AAA-SMC Intermediate |
| **15** | AA | C-SMC |

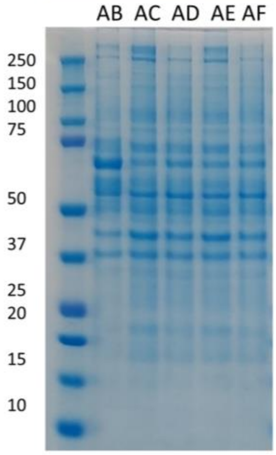


| Slot | Letter | Group |
|---|---|---|
| **1** | **Ladder** | |
| **2** | AB | AAA-SMC Low |
| **3** | AC | AAA-SMC Intermediate |
| **4** | AD | C-SMC |
| **5** | AE | AAA-SMC Intermediate |
| **6** | AF | AAA-SMC Intermediate |

**Figure 1: Coomassie blue staining of samples included in the phosphoproteomics screen for quality control.** Abbreviations: AAA-SMC: Abdominal aortic aneurysm smooth muscle cells, C-SMC: Control smooth muscle cells.

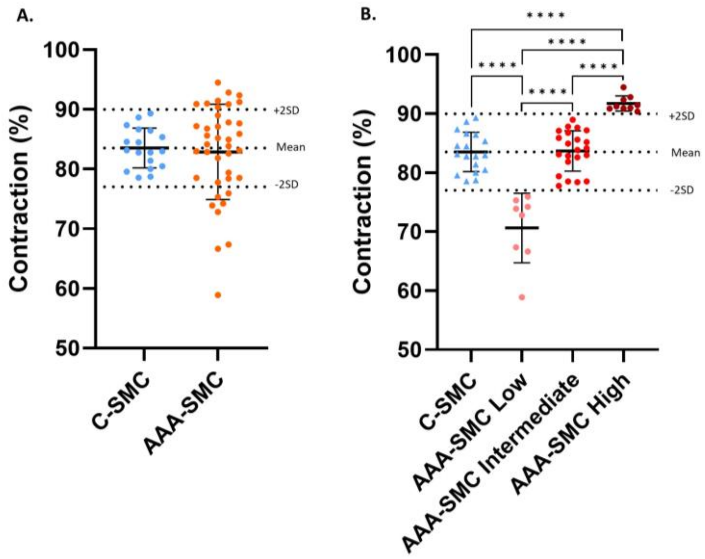


**Figure 2: Contraction measurements in a larger cohort to validate the subdivision of AAA-SMC into three groups based on their contractile response (C-SMC n=18 and AAA-SMC n=39).** Contraction was measured using Electric Cell-substrate Impedance Sensing. The normal vSMC contractility range was defined as the mean contractility of C-SMC, including ±2SD (83,51% ± 6,48%, as indicated by the dotted lines). AAA-SMC were divided into subgroups based on this C-SMC contractility range (AAA-SMC Low contracting n=8, AAA-SMC Intermediate contracting n=22 and AAA-SMC High contracting n=9). The subdivision of the AAA-SMC used in this study into the three contractile groups (as shown in Figure 1B) is similar if these same AAA-SMC would be subdivided based on the ranges calculated in this larger cohort. Cells were seeded in triplicate within each individual experiment and three independent experiments were performed for each vSMC line. Two groups were compared using T-test and multiple groups were compared using one-way ANOVA, with Bonferroni correction. Data is presented as mean with error bars indicating standard deviations. Abbreviations: C-SMC: Control smooth muscle cells, AAA-SMC: Abdominal aortic aneurysm smooth muscle cells, SD: Standard deviation, ****p<0.0001.

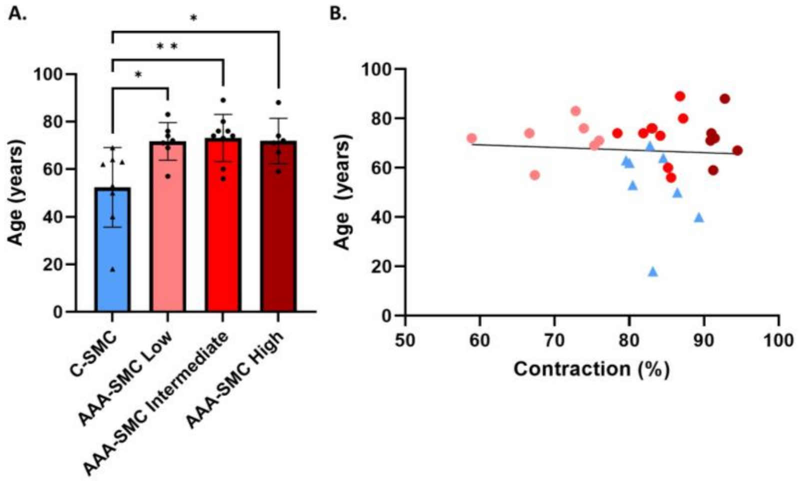


**Figure 3: Age of AAA patients and healthy individuals at the time of the biopsy.** A. Age of the C-SMC (n=8) compared to the three AAA-SMC groups (AAA-SMC Low contracting n=7, AAA-SMC Intermediate contracting n=11 and AAA-SMC High contracting n=6). B. No correlation between age and vSMC contraction. Groups were compared using one-way ANOVA, with Bonferroni correction, and correlation was analyzed with the Pearson correlation. Data is presented as mean with error bars indicating standard deviations. Abbreviations: C-SMC: Control smooth muscle cells, AAA-SMC: Abdominal aortic aneurysm smooth muscle cells, *p<0.05, **p<0.01.

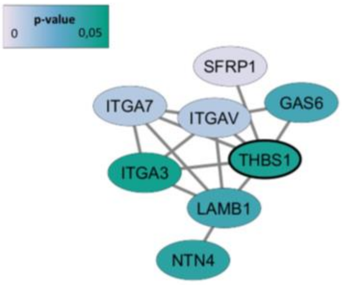


**1. Cell-substrate adhesion (GO ID: 31589)**
8 nodes, p-value: 3.6262^-11

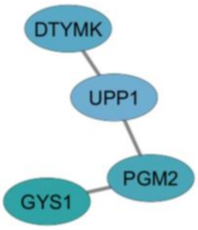


**2. Deoxyribose phosphate metabolic process (GO ID: 19692)**
4 nodes, p-value: 5.9205^-8

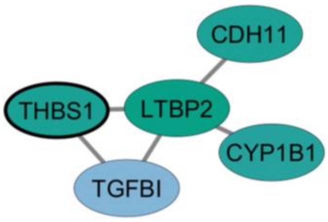


**3. Cell adhesion (GO ID: 7155)**
5 nodes, p-value: 3.8490^-5

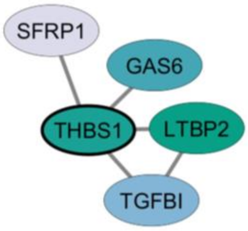


**4. Negative regulation of fibroblast apoptotic processes (GO ID: 2000270)**
5 nodes, p-value: 1.3090^-6

**Figure 4: Cluster and GO analysis on the 60 significantly different expressed proteins between the four groups, as found in the proteomics screen.** Colour of the nodes represent the p-value, as indicated in the figure. Abbreviations: GO: Gene ontology.

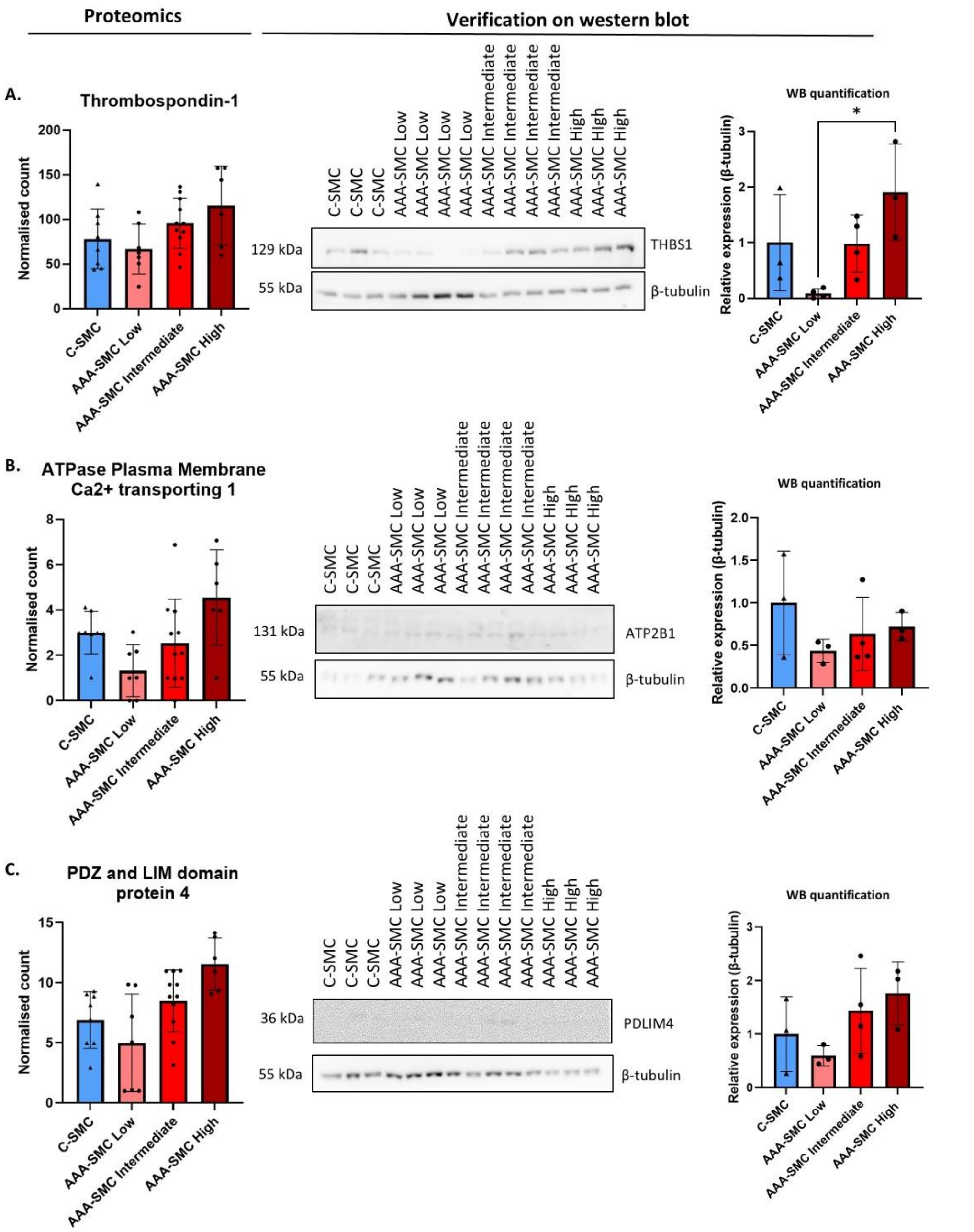


**Figure 5: Verification of the three targets as found in the proteomics screen on WB (C-SMC n=3, AAA-SMC Low contracting n=3/4, AAA-SMC Intermediate contracting n=4 and AAA-SMC High contracting n=3).** A. Verification of Thrombospondin-1 on WB picture and quantification. B. Verification of ATPase plasma membrane $Ca^{2+}$ transporting 1 on WB picture and quantification. C. Verification of PDZ and LIM domain protein 4 on WB picture and quantification. Since the normalized count of ATP2B1 and PDLIM4 as found in the proteomics screen is relatively low, these proteins are challenging to quantify on WB. Groups were compared using one-way ANOVA or Kruskal-Wallis test when not normally distributed, with Bonferroni correction. Data is presented as mean with error bars indicating standard deviations. Abbreviations: C-SMC: Control smooth muscle cells, AAA-SMC: Abdominal aortic aneurysm smooth muscle cells, THBS1: Thrombospondin-1, PDLIM4: PDZ and LIM domain protein 4, ATP2B1: ATPase Plasma Membrane $Ca^{2+}$ transporting 1, WB: Western blot,*$p<0.05$.

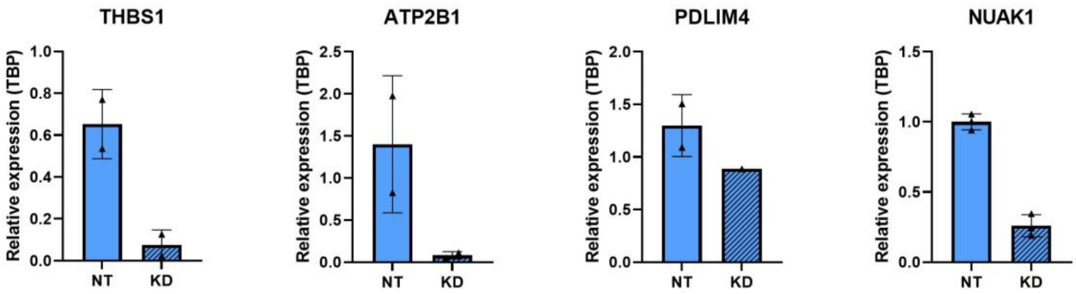


**Figure 6: KD verification of the three selected proteomics targets (THBS1, ATP2B1, PDLIM4) and NUAK1 on RNA level in C-SMC (n=2).** Gene expression was measured using qPCR. Data is presented as mean with error bars indicating standard deviations. Abbreviations: C-SMC: Control smooth muscle cells, KD: Knock down, NT: Non targeting siRNA, THBS1: Thrombospondin-1, PDLIM4: PDZ and LIM domain protein 4, ATP2B1: ATPase Plasma Membrane $Ca^{2+}$ transporting 1, NUAK1: Nu (novel) AMPK related kinase, TBP: TATA-Box Binding Protein, qPCR: quantitative PCR.

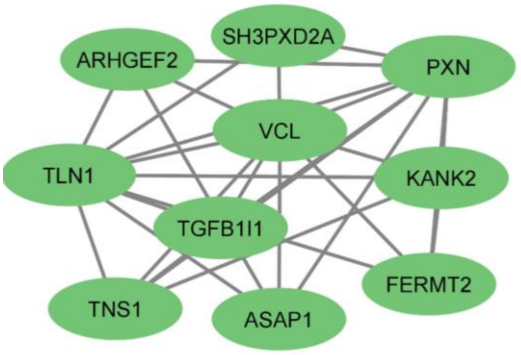


**1. Integrin activation (GO ID: 33622)**
10 nodes, p-value: 1.2599^-5

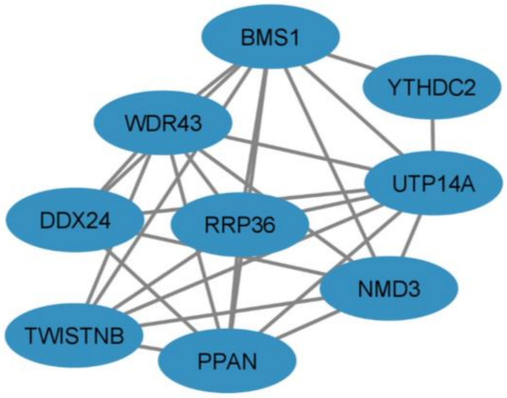


**2. Ribosome biogenesis (GO ID: 42254)**
9 nodes, p-value: 1.4999^-9

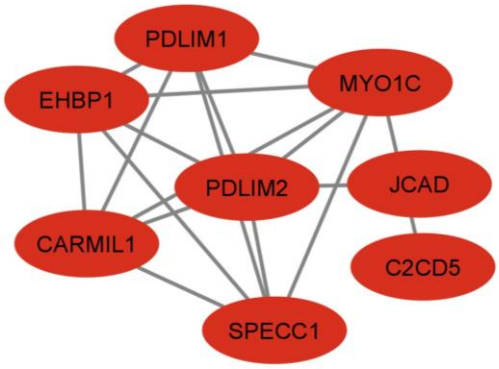


**3. Actin cytoskeleton organization (GO ID: 30036)**
8 nodes, p-value: 2.0115^-8

**Figure 7: Cluster and GO analysis on the proteins belonging to the 396 significantly different phosphorylated phosphosites between the four groups, as found in the phosphoproteomics screen.** Abbreviations: GO: Gene ontology.

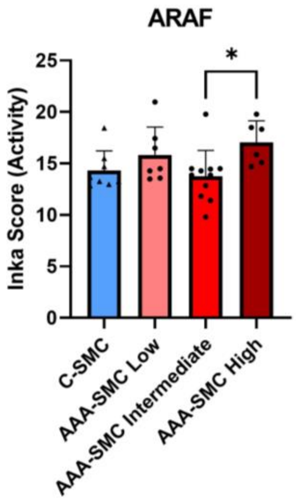


**Figure 8: INKA score of ARAF in the four groups.** INKA score is shown in C-SMC n=8, AAA-SMC Low contracting n=7, AAA-SMC Intermediate contracting n=11 and AAA-SMC High contracting n=6**.** Groups were compared using one-way ANOVA, with Bonferroni correction. Data is presented as mean with error bars indicating standard deviations. Abbreviations: C-SMC: Control smooth muscle cells, AAA-SMC: Abdominal aortic aneurysm smooth muscle cells, ARAF: A-Raf Proto-Oncogene, Serine/Threonine Kinase, INKA: Integrative Inferred Kinase Activity, *p<0.05.

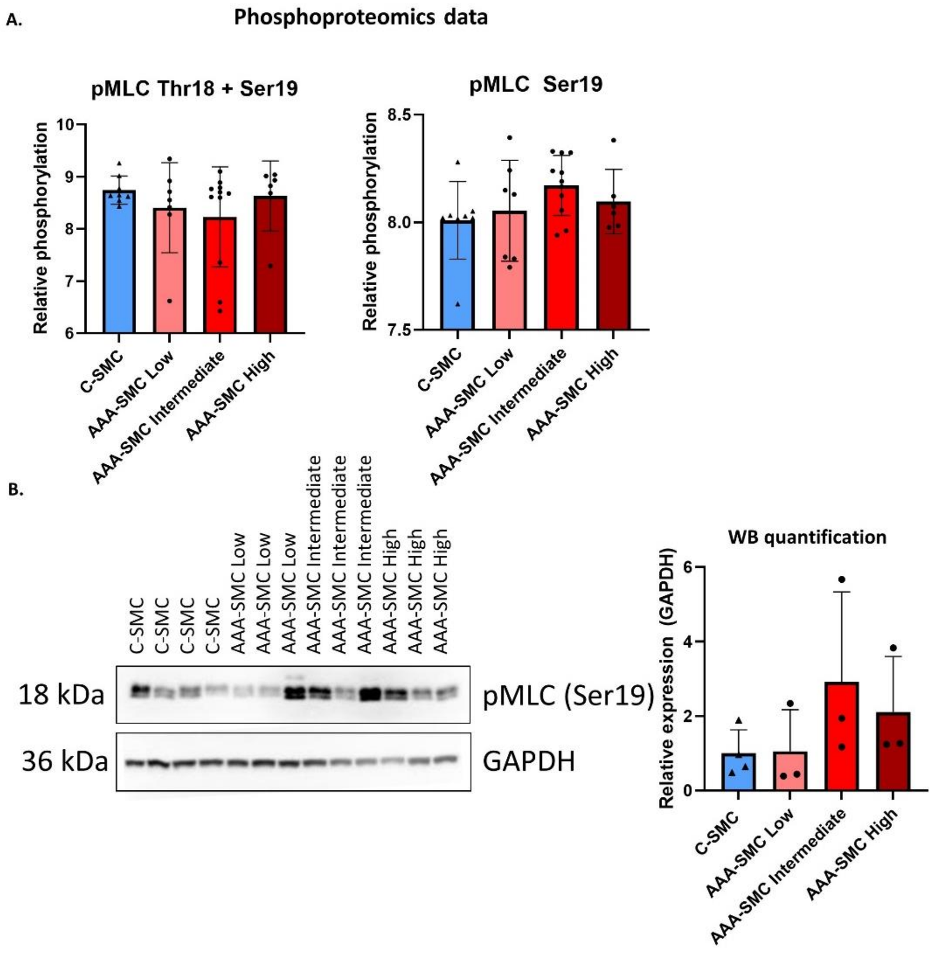


**Figure 9: pMLC levels as found in the phosphoproteomics samples and verified on WB.** A. pMLC levels (left; phosphopeptides with phosphorylated Thr18 and Ser19, right; phosphopeptide with only phosphorylated Ser19) as found in the phosphoproteomics screen in C-SMC n=8, AAA-SMC Low contracting n=7, AAA-SMC Intermediate contracting n=11 and AAA-SMC High contracting n=6. B. pMLC levels (Ser19) on WB picture and quantification in C-SMC n=4, AAA-SMC Low contracting n=3, AAA-SMC Intermediate contracting n=3 and AAA-SMC High contracting n=3. Groups were compared using one-way ANOVA, with Bonferroni correction. Data is presented as mean with error bars indicating standard deviations. Abbreviations: C-SMC: Control smooth muscle cells, AAA-SMC: Abdominal aortic aneurysm smooth muscle cells, pMLC: Phosphorylation of Myosin light chain, GAPDH: Glyceraldehyde-3-Phosphate Dehydrogenase, Ser: Serine, Thr: Threonine, WB: Western blot.

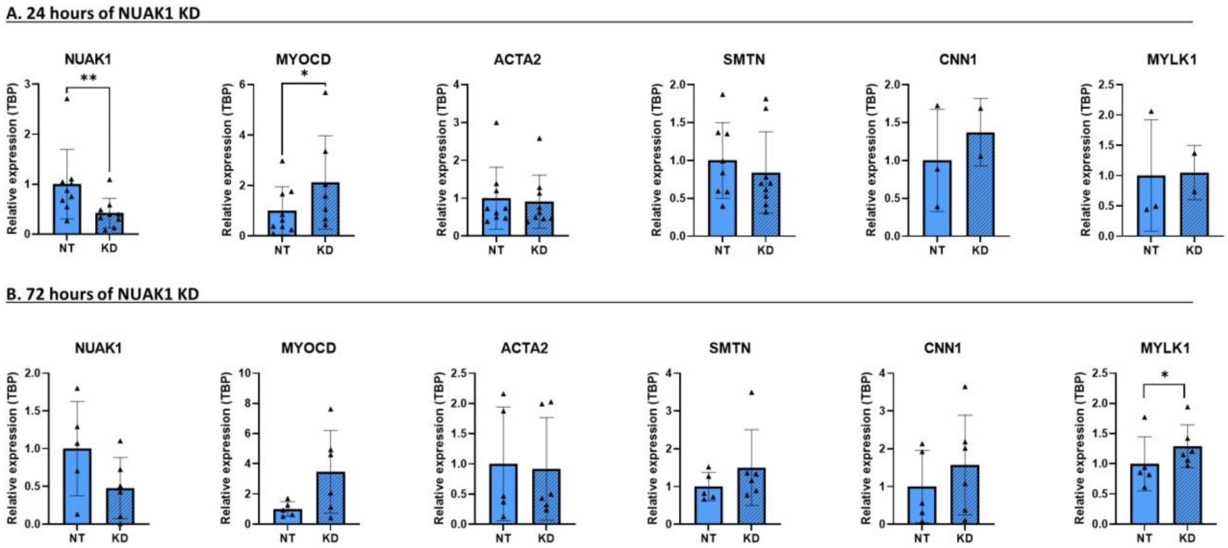


**Figure 10: Effect of NUAK1 KD on gene expression of MYOCD and vSMC contractile markers in C-SMC.** A. Effects of 24 hours of NUAK1 KD on genes encoding for vSMC contractile markers (n=12 or n=3). B. Effects of 72 hours of NUAK1 KD on genes encoding for vSMC contractile markers (n=6). Gene expression was measured using qPCR. Two groups were compared using paired T-test. Data is presented as mean with error bars indicating standard deviations. Abbreviations: C-SMC: Control smooth muscle cells, KD: Knock down, NT: Non targeting siRNA, NUAK1: Nu (novel) AMPK related kinase, MYOCD: Myocardin, ACTA2: Smooth muscle α-actin, SMTN: Smoothelin, CNN1: Calponin, MYLK1: Myosin light chain kinase 1, TBP: TATA-Box Binding Protein, qPCR: quantitative PCR, $^{*}p<0.05$, $^{**}p<0.01$.

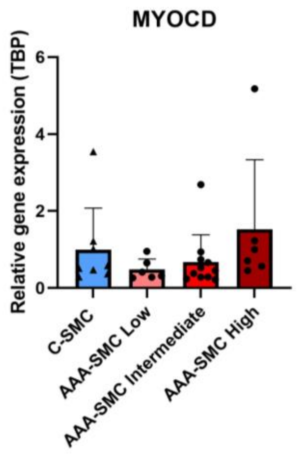


**Figure 11: MYOCD gene expression measured in the same cells as used for the phosphoproteomics screen.** Gene expression was measured using qPCR in C-SMC n=8, AAA-SMC Low contracting n=7, AAA-SMC Intermediate contracting n=11 and AAA-SMC High contracting n=6. Groups were compared using one-way ANOVA, with Bonferroni correction. Data is presented as mean with error bars indicating standard deviations. Abbreviations: C-SMC: Control smooth muscle cells, AAA-SMC: Abdominal aortic aneurysm smooth muscle cells, MYOCD: Myocardin, TBP: TATA-Box Binding Protein, qPCR: quantitative PCR.

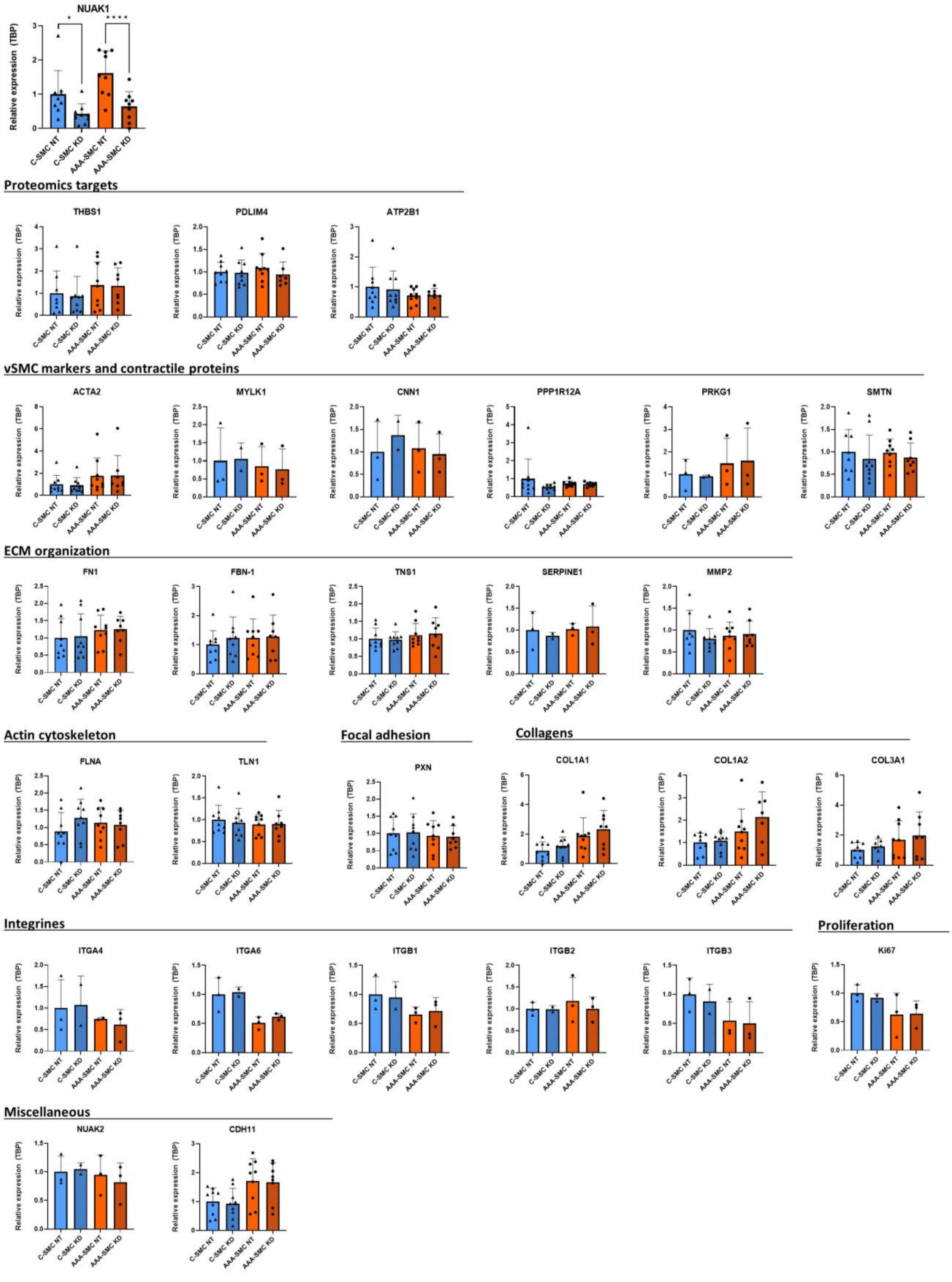


**Figure 12: Effects of NUAK1 KD in C-SMC and AAA-SMC on various genes.** Gene expression was measured using qPCR, in either a cohort of C-SMC n=3 and AAA-SMC n=3, or cohort of C-

SMC n=12 and AAA-SMC n=12. The measured genes encode for the  proteomics targets (THBS1, PDLIM4, ATP2B1), vSMC markers and contractile proteins (ACTA2, MYLK1, CNN1, PPP1R12A, PRKG1, SMTN), ECM organization proteins (FN1, FBN-1, TNS1, SERPINE1, MMP2), actin cytoskeleton (FLNA, TLN1), focal adhesion (PXN), collagens (COL1A1, COL1A2, COL3A1), integrines (ITGA4, ITGA6, ITGB1, ITGB2, ITGAB3), proliferation (KI67) and miscellaneous (NUAK2, CDH11). Two groups were compared using paired T-test. Data is presented as mean with error bars indicating standard deviations. Abbreviations: C-SMC: Control smooth muscle cells, AAA-SMC: Abdominal aortic aneurysm smooth muscle cells, KD: Knock down, NT: Non targeting siRNA, TBP: TATA-Box Binding Protein, qPCR: quantitative PCR, *$p<0.05$, ****$p<0.0001$.
